# Electrochemical carbon fiber-based technique for simultaneous recordings of brain tissue PO_2_, pH, and extracellular field potentials

**DOI:** 10.1101/2020.01.06.895813

**Authors:** Patrick S. Hosford, Jack A. Wells, Isabel N. Christie, Mark Lythgoe, Julian Millar, Alexander V. Gourine

## Abstract

A method for simultaneous electrochemical detection of brain tissue PO_2_ (P_t_O_2_) and pH changes together with neuronal activity using a modified form of fast cyclic voltammetry with carbon fiber electrodes is described. This technique has been developed for *in vivo* applications and recordings from discrete brain nuclei in experimental animals. The small size of the carbon fiber electrode (⍰7μm, length <100μm) ensures minimal disruption of the brain tissue and allows recordings from small brain areas. Sample rate (up to 4 Hz) is sufficient to resolve rapid changes in P_t_O_2_ and pH that follow changes in neuronal activity and metabolism. Rapid switching between current and voltage recordings allows combined electrochemical detection and monitoring of extracellular action potentials. For simultaneous electrochemical detection of P_t_O_2_ and pH, two consecutive trapezoidal voltage ramps are applied with double differential-subtraction of the background current. This enables changes in current caused by protons and oxygen to be detected separately with minimal interference between the two. The profile of P_t_O_2_ changes evoked by increases in local neuronal activity recorded using the described technique was similar to that of blood oxygen level dependent responses recorded using fMRI. This voltammetric technique can be combined with fMRI and brain vessel imaging to study the metabolic mechanisms underlying neurovascular coupling response with much greater spatial and temporal resolution than is currently possible.

## Introduction

The method described here has been developed to study the mechanisms underlying the neurovascular coupling response (Hosford and Gourine 2018). The mechanisms of neurovascular coupling contribute to accurate matching of brain oxygen and glucose supply with demand. Sustained disruption of this balance has been postulated to contribute to cognitive impairment and the development of neurodegenerative disease (Iadecola 2017). Understanding the cellular and molecular mechanisms of neurovascular coupling could be important for the development of future treatments for these conditions. Yet, despite intense experimental scrutiny over the last two decades, the mechanisms underlying neurovascular coupling are not fully understood and are surrounded by controversies (Hosford and Gourine 2018).

The ‘feed-forward’ hypothesis of neurovascular coupling (Attwell et al. 2010) suggests that neurotransmitters released during neuronal activity signal to astrocytes and pericytes to induce dilation of the cerebral vasculature (Mishra et al. 2016). However, astrocytes are also sensitive to changes in partial pressure of oxygen (PO_2_) (Angelova et al. 2015) as well as CO_2_ and protons (H^+^) (Gourine et al. 2010; Howarth et al. 2017; Karagiannis et al. 2016). Changes in brain tissue PO_2_, PCO_2_ and pH correlate with changes in neuronal activity and could contribute to neurovascular coupling via a metabolic feed-back mechanism, as was originally proposed by Roy and Sherrington (Roy and Sherrington 1890).

Blood oxygen level dependent functional magnetic resonance imaging (BOLD-fMRI) (Buxton and Frank 1997) and 2-photon excitation brain vessel imaging (Takano et al. 2006) have been widely used to study the mechanisms of the neurovascular response. However, these techniques have significant limitations. fMRI is non-invasive but suffers from relatively poor spatial and temporal resolution. This is particularly problematic in studies of the brain using small rodent models. Moreover, the BOLD fMRI signal represents a composite response determined by changes in blood flow, blood volume and oxygen consumption, and as such lacks specificity to the underlying haemodynamic mechanisms that give rise to functional hyperaemia. Direct and simultaneous recordings of neuronal activity, although notably achieved during fMRI in non-human primates by Logothetis and co-workers (Logothetis et al. 2001), are not routine. Optical brain imaging is usually confined to cortical structures with imaging depth limited by the light scattering properties of the tissue (Helmchen and Denk 2005). Additionally, to achieve simultaneous recordings of the vessel diameter and neuronal activity one must introduce calcium, voltage sensitive and/or cell specific dyes or genetically encoded sensors of activity. These come with their own caveats; for example there is evidence that commonly used calcium sensors also buffer intracellular calcium and may impair normal cellular function (Bootman et al. 2018). To overcome some of these issues we aimed to develop a minimally invasive technique for *in vivo* recordings of neuronal activity and associated metabolic changes with high spatial and temporal resolution.

Fast-cyclic voltammetry (FCV) using carbon fiber microelectrodes (CFM) is an established technique used to determine the dynamics of neurotransmitter release and reuptake *in vitro* and *in vivo* (Rodeberg et al. 2017). FCV relies on rapidly changing the potential of the CFM vs an Ag/AgCl_2_ reference electrode several times a second over a narrow voltage range to oxidise or reduce the analyte of interest. The amplitude of the current flux between the CFM and the analyte is recorded and used to determine the analyte concentration. Specific advantages of this technique include the small electrode size making it ideal for studies of localised metabolic changes in small brain areas. There is minimal damage to the microcirculation within the tissue and the measurements can be confined to localised brain regions located at any depth. The sub-second time resolution of FCV allows detection of events that occur at the onset of neuronal activity and precede increases in blood flow (Logothetis et al. 2001). Separate electrochemical detection of tissue partial pressure of oxygen (PtO_2_) and pH using FCV has been achieved previously (Bucher et al. 2014; Hosford et al. 2017; Takmakov et al. 2010). Here we describe a novel FCV-based technique that enables simultaneous recordings of key variables representing the metabolic state of the brain: brain tissue PO_2_, pH and neuronal activity.

## Materials and Methods

### Carbon fiber microelectrodes

CFMs (diameter 7 μm) were made as described in detail previously (Hosford et al. 2015; Millar and Pelling 2001). Briefly, single carbon fiber monofilaments (Goodfellow Metals) of ~10cm in length were inserted into borosilicate glass tubing (1.5 mm O.D, Harvard Bioscience) pre-filled with acetone. After complete evaporation of the solvent, the glass tubing was transferred to a conventional horizontal micropipette puller (Model 97, Sutter Instruments) and heated to taper the glass under medium-fast pull speed. The carbon fiber bridging the two pulled electrodes was then cut and connected to a copper wire using a low-melting point tin-bismuth alloy. The carbon fiber tip was trimmed to ~80 μM in length by applying a high DC voltage source (~400 V) guided using a standard laboratory microscope.

### Electrode Calibration

All the recordings were performed using the equipment detailed in Figure 1A. CFMs oxygen sensitivity calibration was performed in phosphate-buffered saline (PBS) containing (in mM) 137 NaCl, 2.7 KCl and 10 phosphate buffer (pH 7.4, unless otherwise stated) saturated with nitrogen to displace the dissolved oxygen. PO_2_ of the solution was then increased stepwise by additions of PBS saturated with 100% oxygen and monitored using an optical oxygen sensor (Oxylite™; Oxford Optronix), adjusted for temperature. A calibration curve of electrode faradaic current changes vs PO_2_ over a range of 5-75 mmHg was constructed.

pH calibration was performed in PBS adjusted to the desired pH using additions of HCL or NaOH and monitored using a standard pH meter. Electrodes were calibrated in a small volume (3 ml) bath that allowed rapid fluid exchange. From a starting point of pH 7.4, buffer pH was changed stepwise to either 7.6, 7.2 or 7.0 in a random order for a particular electrode. A calibration curve of electrode current changes over a pH range of 7.0-7.6 was constructed.

**Figure 1:**
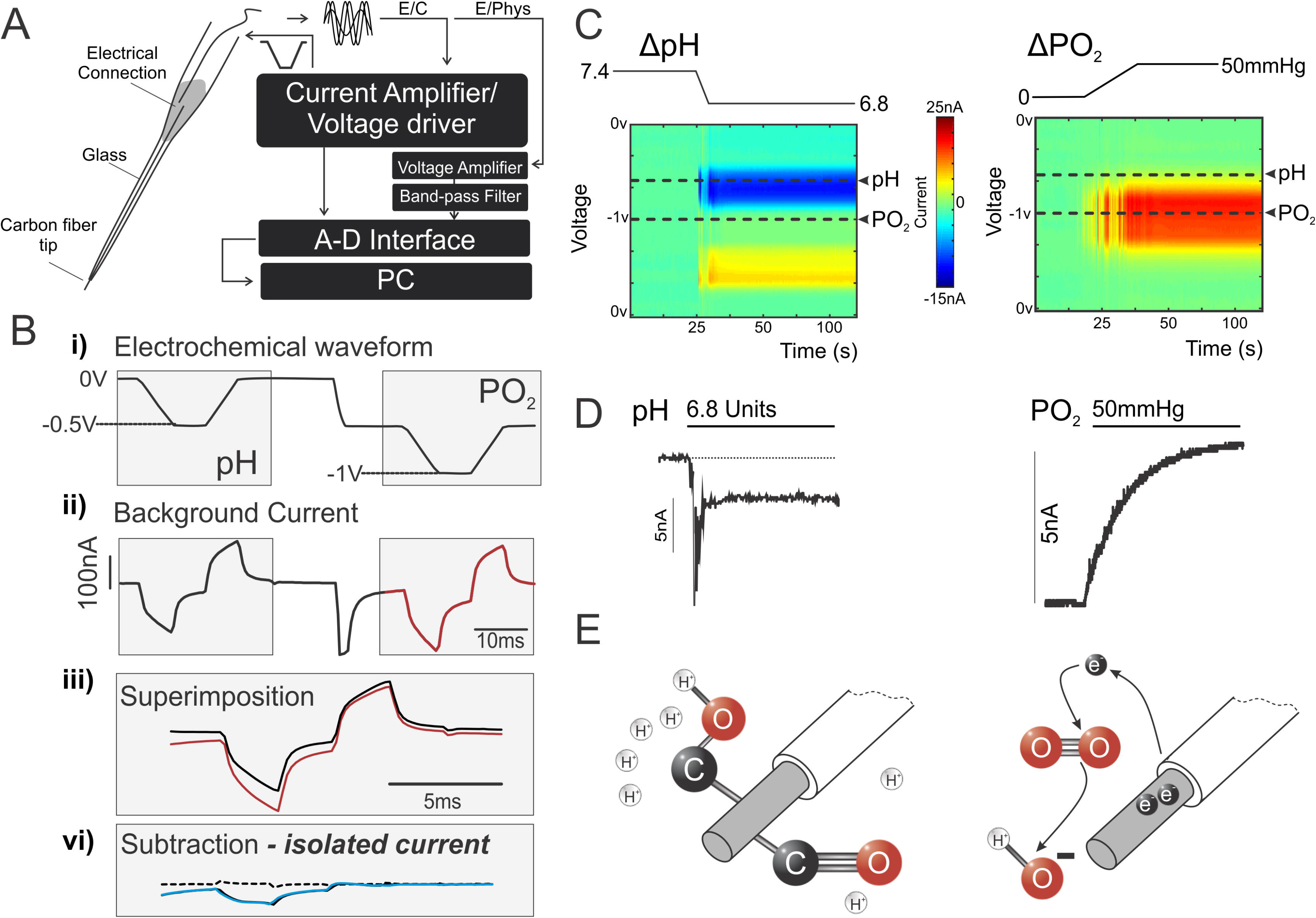
Electrochemical detection of PO_2_ and pH changes using carbon fiber microelectrodes (CFM). **A.** Schematic diagram of the required equipment for simultaneous electrochemical and electrophysiological recordings using CFM. *Left*: construction of carbon-in-glass CFM. *Right*: a block diagram of electrochemical signal generation and acquisition. A voltage driver combined with current amplifier is used for the electrochemical signal generation and detection. A voltage amplifier is used to record neuronal activity between applications of electrochemical waveform. Recorded signal is filtered by a standard band pass filter. All signals are digitised online and recorded for offline analysis. **B.** Electrochemical signal generation and processing: i) two inverted trapezoidal drive voltage waveforms applied to the electrode; the first from 0 to −0.5 V and the second from −0.5 to −1 V. ii) Background current profile generated by the voltage waveform resulting from the capacitive nature of the electrode. iii) Shows the background current resulting from the two applied voltage waveforms combined by superimposition; the resulting combined background current is stored digitally and vi) subtracted from the subsequently acquired scans to provide the measure of the current generated by changes in analyte concentration at the surface of the electrode. **C.** False color plots illustrating current changes over the voltage range of the electrochemical waveform cycling from 0 to −1 V recorded in phosphate-buffered saline. Left shows the current changes induced by a reduction in buffer pH from 7.4 to 6.8 units. *Right* shows the current changes induced by an increase in buffer PO_2_ from 0 to 50 mmHg. Sample points when the peak current changes were measured are indicated for pH and PO_2_. **D.** Representative recordings of CFM electrochemical current changes in response to given changes in pH and PO_2_. **E.** Proposed detection schemes for protons (*left*) and oxygen (*right*) during the voltammetric scan on the surface of the CFM. Protonation of quinone groups on the surface of the CFM produces the current associated with the pH changes. Oxygen is reduced via intermediate steps to hydroxide. The flux of electrons from the CFM to molecular oxygen produces the current associated with changes in PO_2_.

### Ethical approval and animal husbandry

All animal experiments were performed in accordance with the European Commission Directive 2010/63/EU (European Convention for the Protection of Vertebrate Animals used for Experimental and Other Scientific Purposes) and the UK Home Office (Scientific Procedures) Act (1986) with project approval from the University College London Institutional Animal Care and Use Committee. The rats were obtained from Charles River UK and housed in a temperature-controlled room on a 12 h light/dark cycle. Animals had access to standard laboratory chow and water *ad libitum*. On completion of the experiments, the animals were humanely killed by an anaesthetic overdose (pentobarbital sodium, 60 mg kg^−1^, i.v.).

### Animal preparation

Adult male Sprague-Dawley rats (280-320g) were prepared for the experiments in accord with the previously established imaging protocols (Wells et al. 2015). Anaesthesia was induced by isoflurane (2.5-4.0% in oxygen-enriched air) to establish vascular access by femoral vein cannulation. Anaesthesia was then transitioned to α‐chloralose (75 mg kg^−1^, i.v. initial dose followed by supplementary doses of 10–20 mg kg^−1^, i.v., as required). The right femoral artery was cannulated to monitor the arterial blood pressure. The trachea was cannulated and the animal was mechanically ventilated (~60 strokes min^−1^ stroke volume - 8 ml kg^−1^) with oxygen‐enriched room air using a positive pressure ventilator (Harvard Apparatus) or an MR-compatible small animal ventilator (CWE).

Arterial PO_2_, PCO_2_ and pH were recorded at regular intervals using a pH/blood gas analyser (Siemens) and maintained within the physiological ranges (PO_2_ 100–120 mmHg, CO_2_ 35–40 mmHg and pH at 7.35–7.45) by adjusting the frequency and/or volume of mechanical ventilation. Body temperature was maintained at 37.0±0.5°C

### Fast scan cyclic voltammetry

After exposure of the skull surface by midline incision, access to the somatosensory cortex was established via a small craniotomy (~1 mm^2^) The dura was pierced and reflected laterally to prevent damage to the microelectrode tip. Under the microscopic guidance and control of a micromanipulator, CFM was inserted into the somatosensory forelimb region of the cortex (S1FL; coordinates: 2.8-3.8 mm lateral, 1.3-1.5 caudal and 0.1-0.5 mm ventral from Bregma).

The CFM was slowly advanced and placed into the S1FL when clear evoked action potentials were recorded in response to the electrical stimulation (1 Hz) of the contralateral paw. The trapezoidal voltage ramps (Figure 1B) were applied to the CFM (2-4 Hz and 200 Vs^−1^ rate of voltage change.) The CFM signals were amplified (10x), passed through a 50 Hz noise eliminator (Digitimer), filtered to 500-5,000 Hz, digitised (Power1401; Cambridge Electronic Design) and recorded for offline isolation of faradaic currents corresponding to changes in [H^+^] and P_t_O_2_. Continuous switching between current and voltage recordings allowed near-simultaneous detection of the evoked potentials (voltage), [H^+^] and P_t_O_2_ changes (current). Electrical forelimb simulation was applied using a constant‐current stimulator (Digitimer). Trains of stimulation (3 Hz, 1.5 mA, 300 μs pulse width) were applied 3 times per animal/experimental condition with intervals of at least 3 min between the stimulations. Neuronal responses were analysed by integration of the evoked volley of extracellular potentials with the baseline noise subtracted.

### Brainstem recordings

Animals were anaesthetised and instrumented as described above (Figure 1A). In this set of experiments the dorsal surface of the brainstem was exposed for CFM recordings as previously described (Hosford et al. 2015; Hosford et al. 2017). The left cervical vagus nerve was exposed, separated from the sympathetic trunk and placed on the bipolar silver wire electrodes for electrical stimulation (800 μA, 1 ms at 3 Hz).

### fMRI

fMRI was performed using a 9.4T Agilent horizontal bore scanner (Agilent) as described in detail previously (Wells et al. 2015). Briefly, a 72 mm inner diameter volume coil was used for transmission and signal was received using a 4-channel array head coil (Rapid Biomedical). To assess T2* weighted BOLD signals the following sequence parameters were used: TR◻=◻5s, TI◻=◻2s, matrix size◻=◻64◻×◻64, FOV◻=◻35mm × 35mm, TE◻=◻10ms, single slice (slice thickness◻=◻2mm), inversion pulse bandwidth◻=◻20,000Hz (Hosford et al. 2018). BOLD responses in the S1FL region were triggered by electrical stimulation of the contralateral forelimb as described above.

## Results and Discussion

### Voltammetric recordings

Using the recoding setup illustrated in Figure 1A, two negatively-directed trapezoidal voltage ramps were applied to the CFM at an interval of 20 ms as illustrated in Figure 1Bi: the first from 0 to −0.5 V, the second from −0.5 to −1.0 V. Both voltage ramps generated a ‘background’ current due to the impedance of the electrode/ fluid interface (Figure 1Bii). The CFM currents generated during the ramps were digitised and the first was digitally subtracted from the second (Figure 1Biii). This produced a differential background current, (Figure 1Bvi) resulting from the difference between the currents on scans 1 and 2. This subtraction procedure was used to eliminate potential non-specific signals due to changes in tissue impedance, temperature, etc. These non-specific signals appear equally on both the scans and thus are removed by subtraction, allowing discrimination of changes that occur selectively on one or the other scan. Trapezoidal ramps were applied as the background current is predominantly capacitive, reducing this current to a low level during the flat part of the trapezoid where dV/dt is zero.

Using these recording parameters the electrode was found to generate a distinct current profile in response to changes in buffer pH or PO_2_ (Figure 1C). Changes in [H^+^] caused changes in the CFM faradaic current in the voltage range between −0.1 to −0.3 V. These changes occurred only in the first of the two voltammetric scans and, therefore, were clearly present on the differential signal.

Oxygen is electrochemically reduced at a voltage between −0.5 and −1.0 V, generating a cathodal faradaic current. The current from oxygen reduction appears in the signal from the second (−0.5 to −1.0 V) ramp but not from the first ramp and thus could also be seen in the differential signal. Examples of the faradaic current changes in response to oxygen and pH changes in the buffer are illustrated in Figure 1C. There is a clear separation between the two peak currents allowing detection of changes in both analytes simultaneously with minimal interference between the two. Peak current changes are shown in Figure 1D.

### Proposed detection mechanism

The proposed origin of the recorded faradic current is the pH dependent oxidation of the hydroquinone groups on the surface of the CFM (Runnels et al. 1999; Takmakov et al. 2010; Figure 1E). Our recordings support this hypothesis as distinct double-peaks of approximately the same voltage range are detected (Figure 1C). A third pH-dependent peak was reported by Takmakov and colleagues (Takmakov et al. 2010) and was ascribed to changes in electrode capacitance induced by protons disrupting the Helmholtz layer of charged water molecules surrounding the electrode tip and is, therefore, non-faradic in nature. Using double-differential waveform applied during sampling we were able to remove this effect of capacitance change (as it is present on both the waveforms) after subtraction of the resulting background current.

The reaction underlying oxygen detection is the reduction of oxygen to hydroxyl ions (Figure 1E). This reaction occurs in a series of steps involving the formation of intermediates, including hydrogen peroxide. It is non-reversible, as evident from a single unidirectional current peak on the voltammetry scan. There have been previous proposals for the mechanism of electrochemical reduction of oxygen on carbon surfaces (Taylor and Humffray 1975; Zimmerman and Wightman 1991). We propose the following mechanism (Scheme 1) which includes the formation of a hydroperoxyl intermediate as well as hydrogen peroxide.

### CFM calibration

Relative CFM sensitivities to changes in pH and PO_2_ were determined by construction of standard calibration curves in the expected physiological ranges of changes in both variables: 7.0-7.6 units of pH and 5-75 mmHg for PO_2_. Peak faradaic current for pH detection was sampled at −200 mV on each of the scans. Current generated by oxygen was sampled once on each scan at 15 ms from the start of the flat phase of the trapezoid as at this time point the background current was minimal.

*in vitro* calibration demonstrated high CFM sensitivity to oxygen (1.1±0.1 nA per 10 mmHg PO_2_; n = 10 electrodes) and protons (9.8 ± 0.8 nA per pH unit; n = 10 electrodes). Responses were linear within the physiological ranges of changes in these variables (PO_2_, R^2^ =0.998; pH, R^2^ =0.981) (Figure 2A,B). Sensitivity to changes in oxygen did not change over the range of physiological pH values, nor the sensitivity to protons at different PO_2_ levels (Figure 2D). Current responses to a 20 mmHg change in PO_2_ were similar over a range of 7.0-7.6 units: 2.6±0.2 nA at pH 7.0, 2.6±0.2 nA at pH 7.2, 2.7±0.2 nA at pH 7.4 and 2.8±0.2 nA at pH 7.6. Current responses to 0.2 unit decreases in pH were similar over a 10-100 mmHg range of PO_2_: 2.1±0.2 nA at 10 mmHg, 2.1±0.3 nA at 50 mmHg and 2.2±0.3 at 100 mmHg. There was minimal interference between the two measurements; PO_2_-sensitive current was altered by a mere 0.003±0.005nA per 0.1 unit pH change, while pH-sensitive current changed by 0.1±0.03 nA per 10 mmHg change in PO_2_ (n =6, Figure 2C and E).

**Figure 2:**
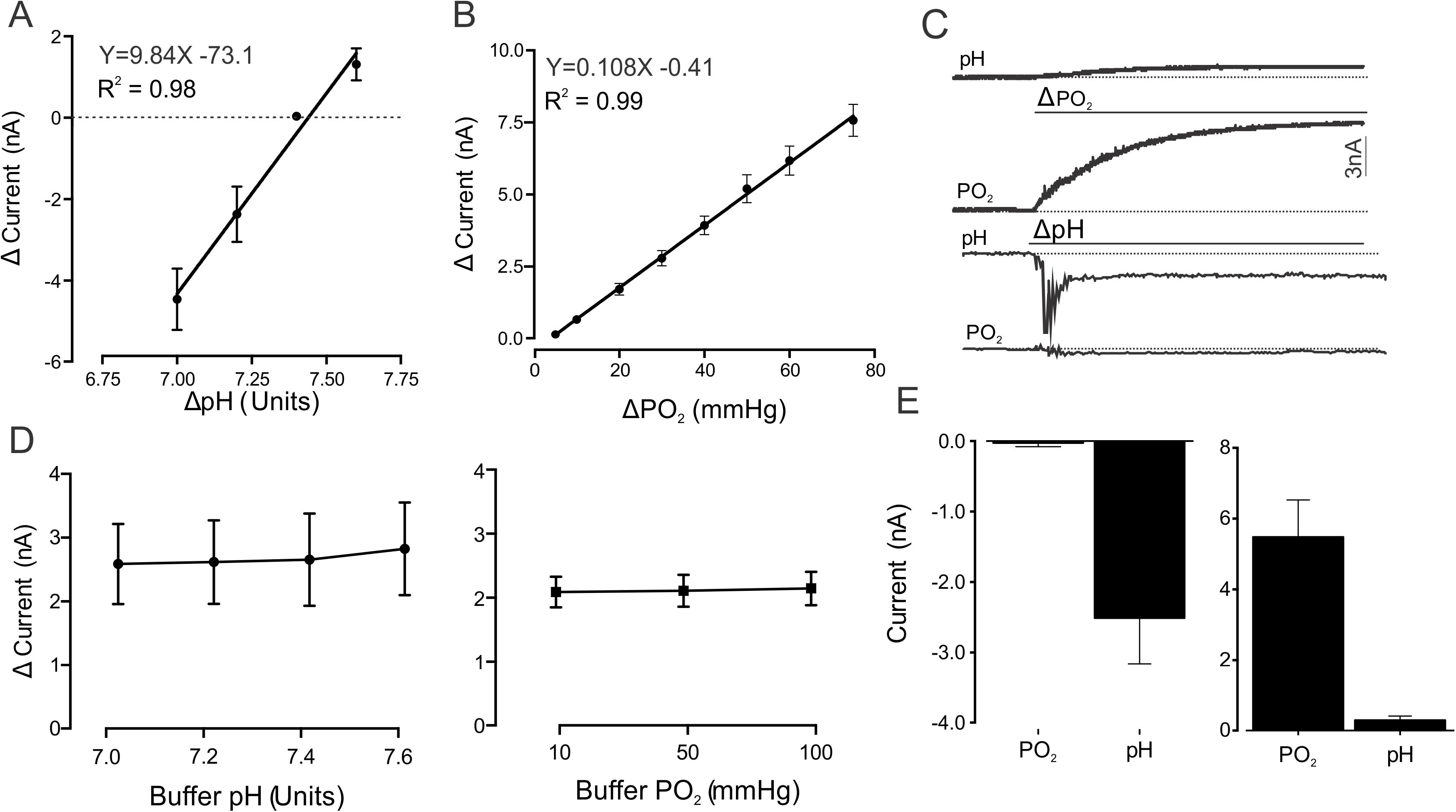
Validation of the technique and calibration of CFMs. **A.** Calibration curve illustrating changes in CFM electrochemical current recorded at the pH sample point in response to changes in buffer pH from 7.4 to 7.0, 7.2 and 7.6. Calibration points represent the means (±S.E.M) peak current changes of 10 electrodes exposed to a given change in buffer pH starting from 7.4. **B.** Calibration curve illustrating changes in CFM electrochemical current recorded at the oxygen sample point in response to changes in buffer PO_2_ between 5-75 mmHg. Calibration points represent the means (±S.E.M) peak current changes of 10 electrodes, each exposed to every PO_2_ increment once. Standard linear regression was applied to fit a line for both calibrations; the equation and R^2^ are indicated. **C.** Representative traces showing changes in peak current at the PO_2_ and pH sample points during a decrease in buffer pH of 0.2 units or increase in buffer PO_2_ by 50 mmHg. **D.** *Left*: summary data showing peak currents generated in response to a 20 mmHg increase in PO_2_ over a physiological range of changes in buffer pH at 7.0, 7.2, 7.4 or 7.6 units. Each point represents the mean (±S.E.M) peak response of 12 electrodes. *Right*: summary data showing peak currents generated in response to a 0.2 unit decrease in pH over a physiological range of changes in PO_2_ at: 10, 50 and 100 mmHg. Each point represents the mean (±S.E.M) peak response of 9 electrodes. **E.** Summary data of peak current changes recorded at the pH and PO_2_ sample points in response to a decrease in pH by 0.2 units (*left*) and at the pH and PO_2_ sample points in response to an increase in PO_2_ by 50 mmHg (*right*). Shown are the means (±S.E.M) peak current responses of 6 electrodes.

These recordings show that there is no significant change in current produced by one analyte when the other varies over the physiological range expected in the brain tissue. Cross-talk between the pH and oxygen detection signals was calculated to be less than 5%, which is within the range reported for other electrochemical detection techniques (Tian et al. 2009). Further, we can confidently exclude potential contamination of the recorded signals by other oxidizable molecules as these require a positive waveform voltage, usually +1.3 V (Park et al. 2011), while this technique records PO_2_ and pH changes using a negative (reducing) waveform of 0 to −1 V.

### *In vivo* application

Using the recording setup illustrated by Figure 3A, changes in brain P_t_O_2_ were recorded during systemic hypoxia induced by a 20s-long suspension of the mechanical ventilation. This resulted in an immediate sharp decrease in oxygen-associated faradaic current by 3.2±0.6 nA, equivalent to a reduction in brain P_t_O_2_ by 19±3 mmHg (n =6; Figure 3Bi and ii). Upon re-instatement of lung ventilation, the brain P_t_O_2_ rapidly reversed and exceeded the baseline level by 15±4 mmHg within 10 s (n =6; Figure 3Bii).

**Figure 3:**
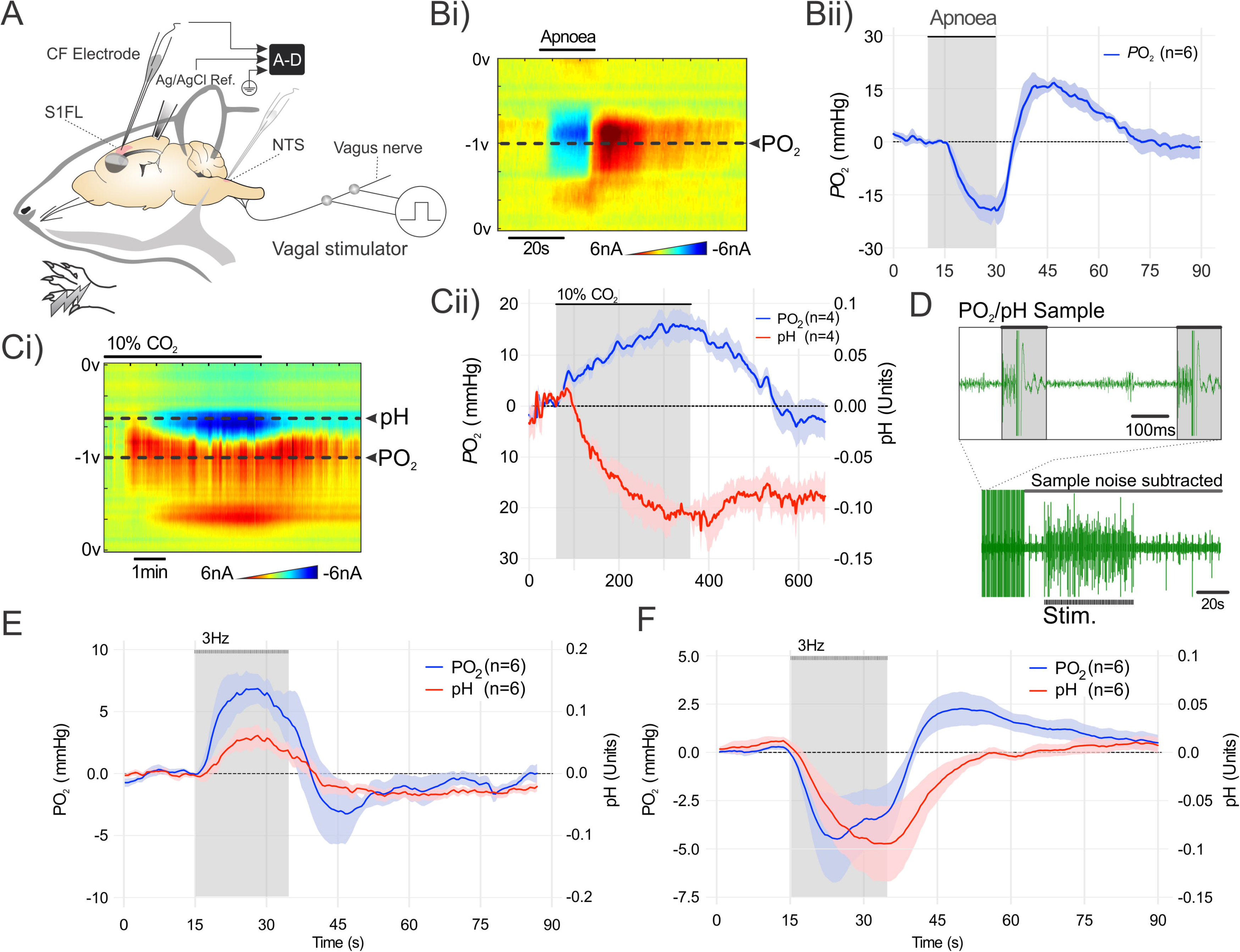
Simultaneous detection of brain tissue PO_2_, pH, and extracellular field potentials. **A.** Schematic illustration of the *in vivo* recording setup in an anesthetised rat showing placement of the CFM in the somatosensory cortex with Ag/AgCl reference electrode in the contralateral hemisphere. The forepaw was stimulated electrically to recruit somatosensory pathways. Separate placement of the CFM in the nucleus of the solitary tract (NTS) and the vagus nerve stimulator are also shown. **B.** The effect of apnoea on oxygen-sensitive current recorded in the somatosensory cortex. i) False color plot showing changes in oxygen-sensitive current over the range of the voltammetric scan during a 20 s apnoeic episode. The sample point from which the oxygen-sensitive current was recorded is indicated. ii) Brain P_t_O_2_ changes recorded using CFM during systemic hypoxia (means±S.E.M; n =6) **C.** The effect of respiratory acidosis induced by inhalation of 10% CO_2_ in the inspired gas mixture on oxygen and proton-sensitive currents recorded in the somatosensory cortex. i) False color plot showing changes in oxygen and proton-sensitive currents over the range of the voltammetric scan during systemic hypercapnia. The sample points from which the oxygen and proton-sensitive currents were recorded are indicated. ii) Brain tissue PO_2_ and pH changes recorded using CFM during systemic hypercapnia (means±S.E.M; n =4) **D.** Representative recording of extracellular field potentials in the somatosensory cortex evoked by electrical forelimb stimulation during simultaneous brain P_t_O_2_ and pH sampling. Inset illustrates the intervals between pH/PO_2_ sampling available for extracellular potential recording. **E.** Changes in brain P_t_O_2_ and pH recorded in the somatosensory cortex in response to electrical forelimb stimulation (3 Hz, 20 s; means ±S.E.M; n =6). **F.** Changes in P_t_O_2_ and pH recorded in the nucleus of the solitary tract of the brainstem in response to electrical stimulation of the vagus nerve (3 Hz, 20 s; means ±S.E.M; n =6).

Changes in brain P_t_O_2_ and pH were next recorded during systemic respiratory (hypercapnic) acidosis induced by CO_2_ inhalation (10% CO_2_ in the inspired gas mixture; 5 min). This stimulus caused a significant decrease in current by 0.31±0.05 nA (Figure 3Ci) in the voltage range corresponding to pH changes, equivalent to a decrease in brain tissue pH by −0.11±0.02 units (n =4; Figure 3Bii). Systemic hypercapnia also caused a 0.54±0.1nA increase in current (n =4; Figure 3Bii) over the voltage range corresponding to changes in PO_2_, equivalent to an increase in brain P_t_O_2_ by 15±3 mmHg. This reflected CO_2_-induced increase in global brain blood flow. Upon return to normocapnia the brain P_t_O_2_ decreased back to baseline within 3 min. Partial recovery of brain tissue pH was recorded during the same time period. The faradic current profiles recorded during systemic hypoxia and CO_2_-induced acidosis were found to correspond closely to similar responses to changes in PO_2_ and pH induced *in vitro*. Following subtraction of the control scan current from the active scan current, increases in oxygen concentration produce positive and increases in proton concentration produce negative current increments.

Electrical stimulation of the forepaw increased the neuronal activity in the S1FL of the cortex as was evident from an increase in action potential firing (recorded by the CFMs during the intervals between the applications of the voltage ramps; Figure 3D). Traces depicted in Figure 3D show the extracellular spike activity in S1FL, time-locked to the application of the stimulus.

Electrical stimulation of the forepaw was associated with a consistent increase in faradic current corresponding to an increase in P_t_O_2_ (n =6; Figure 3E). Increases in P_t_O_2_ were observed 1-2 seconds after the onset of the stimulation. The response was found to be biphasic with an initial increase during the period of stimulation followed by a post-stimulus decrease below the baseline (Figure 3E). Calibration of the CFM after each of the recordings revealed the peak increases in P_t_O2 of 6.9±1.2mmHg and the post-stimulus decreases with the magnitude of 3.2±2.5 mmHg (n =6; Figure 3E). Activation of somatosensory pathways concomitantly increased the faradic current recorded at the sample point for the detection of pH changes. Changes in brain tissue extracellular pH followed a similar time course, but the pH response lagged the PO_2_ changes by ~1 s. The pH signal displayed biphasic response profile with initial alkalisation of 0.06±0.02 pH units during the period of stimulation, followed by a decrease of 0.03±0.01 below baseline after the termination of the stimulus (n =6; Figure 3E).

There is evidence that the mechanisms of neurovascular coupling might be different in different brain areas (Devonshire et al. 2012). We next placed the CFM within the nucleus of the solitary tract (NTS) of the brainstem, and recorded changes in PtO_2_ and pH evoked by activation of *visceral* sensory pathways. The NTS receives mono-synaptic afferent inputs via the vagus nerve (Berthoud and Neuhuber 2000). Electrical stimulation of the vagus nerve produced biphasic changes in both P_t_O_2_ and pH in the NTS that were markedly different from those recorded in the somatosensory cortex. There was an initial decrease in P_t_O_2_ by 4.5±2.2 mmHg and extracellular acidification by 0.095±0.04 pH units, followed by a post-stimulus overshoot in P_t_O_2_ by 2.3±0.9 mmHg (n =6; Figure 3F) with pH slowly recovering towards the baseline.

The P_t_O_2_ and pH recordings performed within the NTS show that this technique is applicable to studies of small discrete nuclei and/or regions located deep in the brain that are difficult to access using the existing imaging techniques. This could be especially useful when investigating the heterogeneity of the neurovascular coupling responses in the brain, as highlighted by dramatically different P_t_O_2_ and pH response profiles recorded in the NTS (Figure 3F) and the somatosensory cortex (Figure 3E).

### Comparison with fMRI

BOLD signals induced in the S1FL by activation of somatosensory pathways were recorded in identical experimental conditions to allow comparison between the responses recorded using fMRI and P_t_O_2_ changes recorded using voltammetry. Electrical forepaw simulation induced biphasic BOLD signal changes in the S1FL (Figure 4A and B). In order to compare the BOLD signal changes with measured changes in P_t_O_2_, the calibrated voltammetry signal was down-sampled to 0.5Hz and both signals were standardised with a z-score function and overlaid (Figure 4C). Plotting the distance between each sample point for the two techniques revealed much of the difference observed during the post-stimulus undershoot, where it was maximal 6 s after the termination of the stimulus.

**Figure 4:**
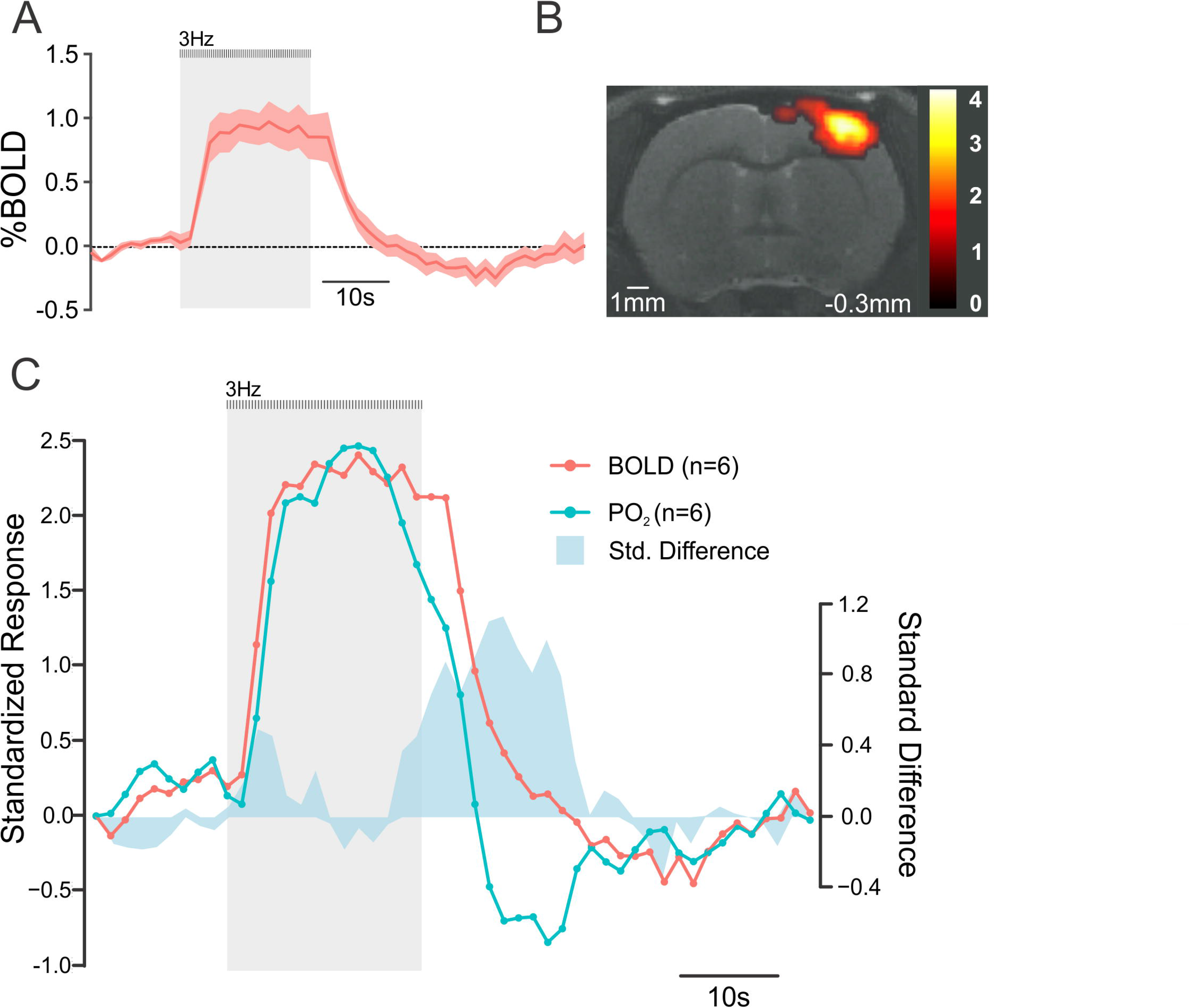
Comparison of brain P_t_O_2_ changes profile recorded using CFM voltammetry and that of BOLD responses recorded using fMRI. **A.** BOLD response profile illustrating changes in mean signal intensity in the somatosensory cortex (S1FL) induced by the electrical stimulation of the contralateral forelimb in anesthetised rats (means ±◻JS.E.M; n =6). **B.** Representative BOLD activation map (familywise error, p◻<◻0.05, nv◻ =◻3) taken at one coronal slice (distance from Bregma is indicated) level showing activation of the somatosensory cortex in an anesthetised rat. Color bar represents the t-score from statistical parametric mapping mixed-effects analysis, p < 0.05 (uncorrected). **C.** Comparison of time course and response profile of standardised P_t_O_2_ and BOLD responses in the S1FL of the anesthetised rat evoked by forelimb simulation. Responses are represented as a mean z-score from the recordings obtained in 6 animals for each experimental measurement. The standard difference is the difference calculated between the two z-scores at each sample point. P_t_O_2_ recordings were down sampled by averaging 8 sampled data points to achieve the sampling frequency of 0.5 Hz, equal to that of the fMRI sampling rate.

The data obtained show that the profile of brain PtO_2_ changes recorded using this voltammetric technique is virtually identical to the profile of BOLD responses recorded using fMRI, with additional advantage of simultaneous detection of brain tissue pH and monitoring of the evoked neuronal activity.

### Conclusion

Here we describe a novel experimental technique for simultaneous detection of brain P_t_O_2_, pH and extracellular field potentials using CFM voltammetry. Electrochemical detection of P_t_O_2_ and pH changes with near-simultaneous recordings of neuronal activity is possible in small nuclei located deep in the brain. Simultaneous monitoring of blood flow (P_t_O_2_), metabolism (pH) and neuronal activity using the CFM-based technique described here may prove to be useful in studies of the metabolic mechanisms underlying the neurovascular coupling response.

## Acknowledgements

The author(s) disclose receipt of the following financial support for the research, authorship, and/or publication of this article:

This work was supported by The Wellcome Trust and British Heart Foundation

JAW is a Wellcome Trust/Royal Society Sir Henry Dale Fellow

AVG is a Wellcome Trust Senior Research Fellow (Refs: 095064 and 200893).

The author(s) declare no potential conflicts of interest with respect to the research, authorship, and/or publication of this article.

---

**Scheme 1:**
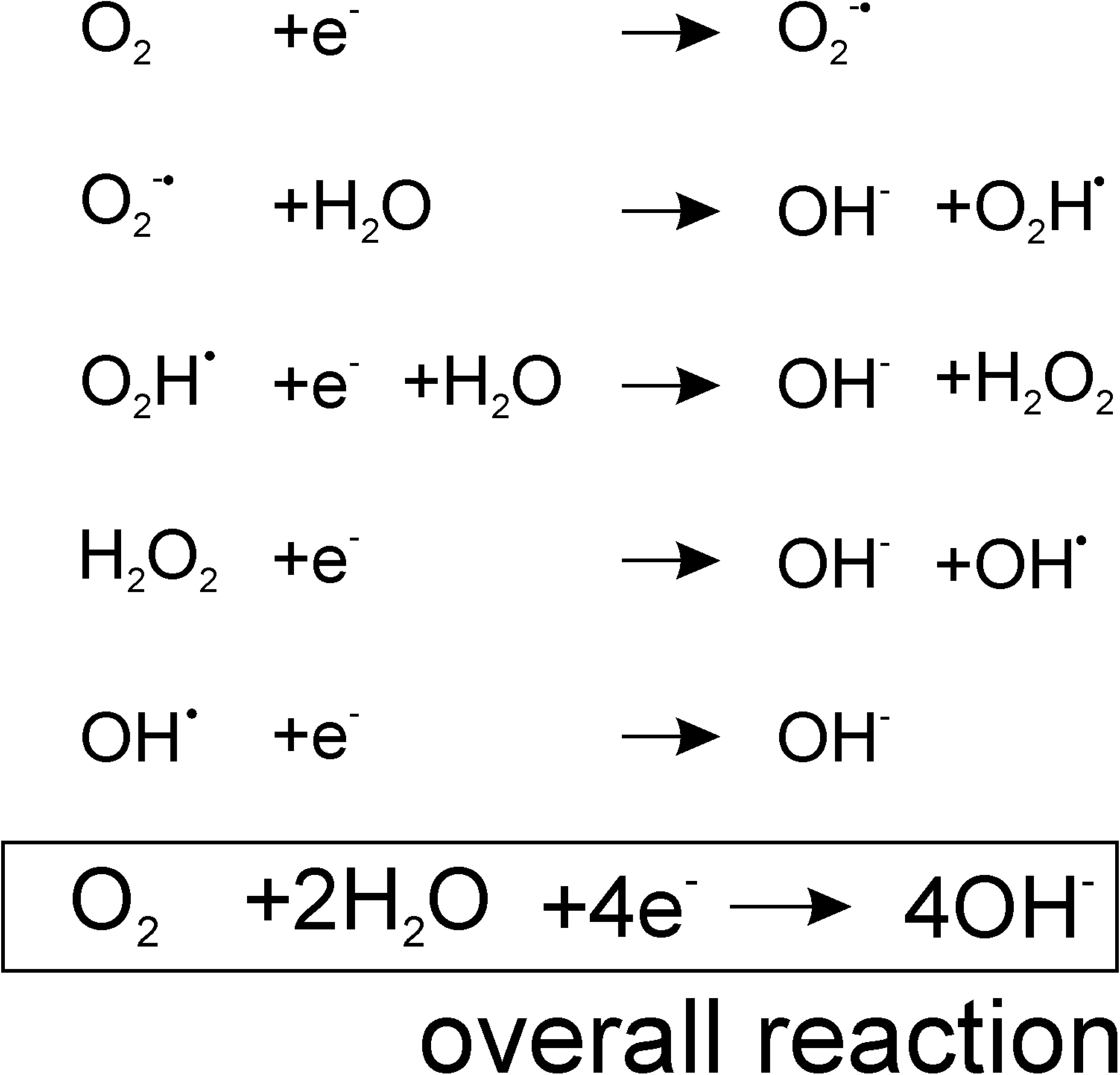
Proposed intermediate and overall reaction scheme underlying the reduction of molecular oxygen on the carbon surface.

